# EnsembleSeq: A workflow towards real-time, rapid and simultaneous multi-kingdom amplicon sequencing for holistic and cost-effective microbiome research at scale

**DOI:** 10.1101/2023.12.09.570917

**Authors:** Sunil Nagpal, Sharmila S. Mande, Harish Hooda, Usha Dutta, Bhupesh Taneja

## Abstract

**Background:** Bacterial communities are often concomitantly present with numerous microorganisms in the human body and other natural environments. Amplicon based microbiome studies have generally paid a skewed attention, that too at a rather shallow genus level resolution, to the highly abundant bacteriome, with interest now forking towards the other microorganisms, particularly fungi. Given the generally sparse abundance of other microbes in the total microbiome, simultaneous sequencing of amplicons targeting multiple microbial kingdoms could be possible even with full multiplexing. Guiding studies are currently needed for performing and monitoring multi-kingdom-amplicon sequencing and data capture at scale.

**Method:** Full length bacterial 16S rRNA gene and entire fungal ITS region amplification was performed for human saliva samples (n=96, including negative and positive controls). Combined amplicon DNA libraries were prepared for nanopore sequencing using a major fraction of 16S molecules and a minor fraction of ITS amplicons. Sequencing was performed in a single run of an R10.4.1 flowcell employing the latest V14 chemistry. An approach for real time monitoring of the species saturation using dynamic rarefaction was designed as a guiding determinant of optimal run time.

**Results:** Real-time saturation monitoring for both bacterial and fungal species enabled the completion of sequencing within 30 hours, utilizing less than 60% of the total nanopores. ∼5 million HQ taxonomically assigned reads were generated (∼4.2 million bacterial and 0.7 million fungal), providing a wider (beyond bacteriome) snapshot of human oral microbiota at species level resolution. Among the more than 400 bacterial and 240 fungal species identified in the studied samples, the species of Streptococcus (e.g. *S. mitis, S. oralis*) and Candida (e.g. *C. albicans, C. tropicalis*) were observed to be the dominating microbes in the oral cavity, respectively. This conformed well with the previous reports of the human oral microbiota.

**Conclusion:** Ensembleseq provides a proof-of-concept towards identification of both fungal and bacterial species simultaneously in a single fully multiplexed nanopore sequencing run in a time and resource effective manner. Details of this workflow are provided to enable large scale application for a holistic species level microbiome study.

## 1. Introduction

Human microbiome represents the total gene pool of bacteria, fungi, viruses, protists and other microscopic organisms inhabiting the human body (1). Bacteria (bacteriome) have traditionally received particular interest, and in fact rather synonymously (mis)represented as the total ‘microbiome’. This is largely due to the historical focus on bacteria since the times of Leeuwenhoek (2), ease of bacterial culturing, an understanding that bacteria are the most abundant microbial cells in human body and a general lack of cost-effective approaches to simultaneously study the total human microbiome (2, 3). The sparse population of other microorganisms, understood to constitute the dark matter of human microbiome largely remain to be (taxonomically and functionally) elucidated (3, 4). Recent reports of the human gut and oral mycobiome, especially the disorder linked mycobiota (e.g., the pan-cancer fungi-typing), have now highlighted the understudied diversity and importance of the rare biosphere of the human body (5–7). Evidence is now accumulating towards not only the diagnostic potential of site-specific fungal communities in the human body but also their cross-talk with the dominating but co-inhabiting bacteriome (5–8). These strides for studying the mycobiome, are encouraging and much needed for moving towards the holistic understanding of the (human) microbiome.

Amplicon sequencing has conventionally offered a cost-effective way of characterizing the human microbiome (9). Until the advent of long-read third generation sequencing techniques (10), bacterial communities were profiled by amplifying and sequencing the short (∼300-500bp) segments pertaining to one or more of the (traversing V1 toV9) variable regions of the ∼1.5kb long 16S rRNA gene (1, 9, 11, 12). The short segments of the variably long internal transcribed spacer region (ITS) region, (spanning ITS1, 5.8S and ITS2) between the 18S and 28S rRNA genes, on similar lines served as universal barcodes for fungi (6, 8, 13, 14). Sequencing of the short segments of the marker genes however constrained the taxonomic classification primarily at the genus level, hence often limiting the majority of the (amplicon based) human microbiota research to a rather under-resolved or shallow profiling (15, 16). Long-read sequencing technology has circumvented these limitations and enabled much resolved (species and even strain level) characterization of microbial communities by targeting the full-length of 16S and ITS regions (15–17). This advantage has reignited a deeper, species level probing of the previously genus-level characterized human microbiota (15, 18, 19). At this juncture of revisiting the microbiome, skewed attention still remains towards the highly abundant bacteriome, with a limited number of independent studies reporting the characterization of human fungal microbiota using the entire ITS or large subunit (LSU) regions (16–18, 20–28).

A holistic view of the microbial community of any environment must include information of the total microbiota (spanning different microbial kingdoms) to probe the role of the multiple prevalent microorganisms and their cross-talks (1, 3, 8). While this is possible through whole metagenome sequencing, the cost of obtaining sufficient depth for a reliable assembly of all microbial genomes, overcoming the noise of rather dominant human or eukaryotic DNA, can be prohibitive, especially for large cohort studies (29, 30). Notably, even the amplicon sequencing can turn expensive if for every target microorganism (or amplicon type) separate sequencing runs (and/or upstream library preparation steps) are performed. Multi-amplicon sequencing (i.e., combining the amplicon DNA from multiple genes or regions), usually employed for increasing the specificity of taxonomic profiling through assembly and consensus of multiple regions, can potentially also provide a cost-effective way of capturing different kingdoms of microorganisms in the same set of samples (12, 31).

Human microbiome is predominantly composed of the bacterial cells (e.g., ∼10^11^-10^12^ CFU/g of cultivable bacteria in the faeces or 10^9^-10^10^ CFU/ml of saliva in the oral cavity) (1, 32). A sparse content of other microorganisms, also called as the rare biosphere, is also observed in the human microflora (e.g., fungi, comparably forming ∼10^3^-10^6^ CFU/g of faeces or 10^1^-10^4^ CFU/ml of saliva) (1, 4, 32, 33). This inherent sparsity of non-bacteriome populations in the total microbiome can perhaps offer an advantage in concomitantly sequencing a fraction of their (few kilobase long) marker amplicons (∼ 0.6 kb ITS region) along with the major proportion of the bacterial amplicons (∼ 1.5 kb 16S gene) without prejudicing the coverage of the either. This approach, here termed as “multi-kingdom amplicon sequencing” may potentially lower the time, effort and cost of microbial community evaluation while also simultaneously characterizing the co-occurring endogenous residents of the human microbiota. A few recent reports, though using short reads and a small number of samples, have in fact provided encouraging evidence towards potential to concomitantly capture both bacteriome and mycobiome using amplicon sequencing (31, 34). Current literature evidence however seems to indicate that a majority of non-bacteriome (primarily mycobiota) studies are done in silos (e.g., only mycobiome profiling of oral cavity) (5, 14, 20, 22, 35, 36) with a handful of short read microbiome studies attempting both bacterial and fungal amplicon sequencing for the targeted samples, often as independent libraries or even independently sequenced bioprojects (31, 33, 37–39). Studies showcasing the success (and even failures) of the multi-kingdom amplicon sequencing approach, with clearly outlined protocols may add to the currently very limited evidence and encourage the (resource constrained) researchers to expand the scope of their microbiome research. This is also particularly important for the long-read technology and multiplexed experiments, which may demand high coverage or impact the sequence sufficiency, guiding case studies for which are lacking.

Here, as part of resource optimization effort for a larger microbiome cohort study in our lab, we present a guided workflow entitled, EnsembleSeq (summarized in Figure 1 and Figure 2) as a strengthening proof-of-concept case study (first using the nanopore technology) for a holistic multi-kingdom human oral-microbiome sequencing. Briefly, following avenues were explored that led to the design of EnsembleSeq: (i) attempting simultaneous (multi-kingdom) sequencing by spiking bacterial amplicons with a fraction of fungal amplicons (ii) analysing whether the spiking sufficiently captures the bacteriome as well as the mycobiome (iii) devising an approach towards guiding the decisions for optimal run-time and timely decisions to avoid overutilization of the reusable flowcells, without impacting species saturation. Furthermore, the concordance of the concomitantly captured human oral bacteriome and mycobiome with the previous reports was also validated.

**Figure 1.**
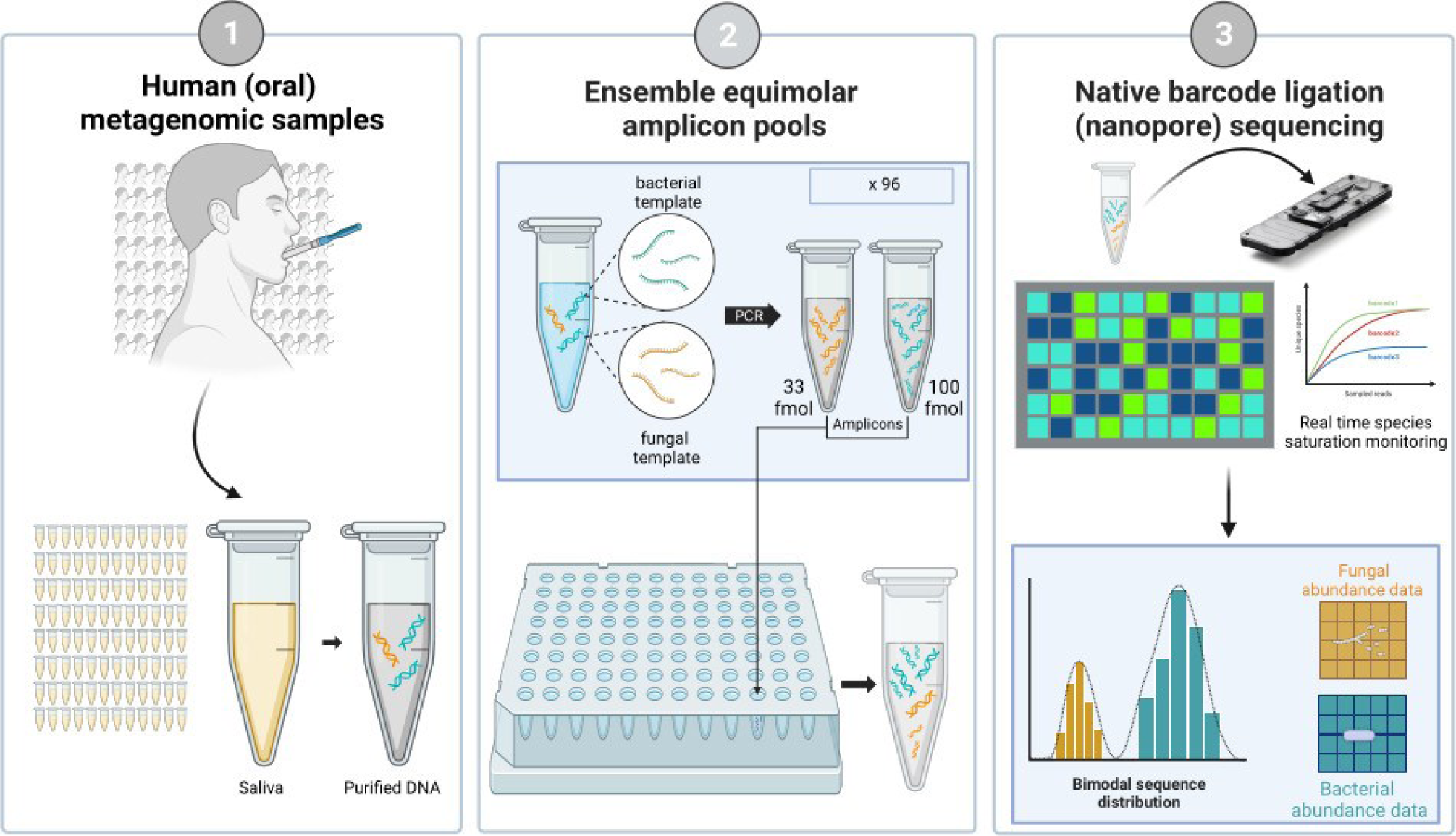
Graphical overview of the design of EmsembleSeq workflow

**Figure 2.**
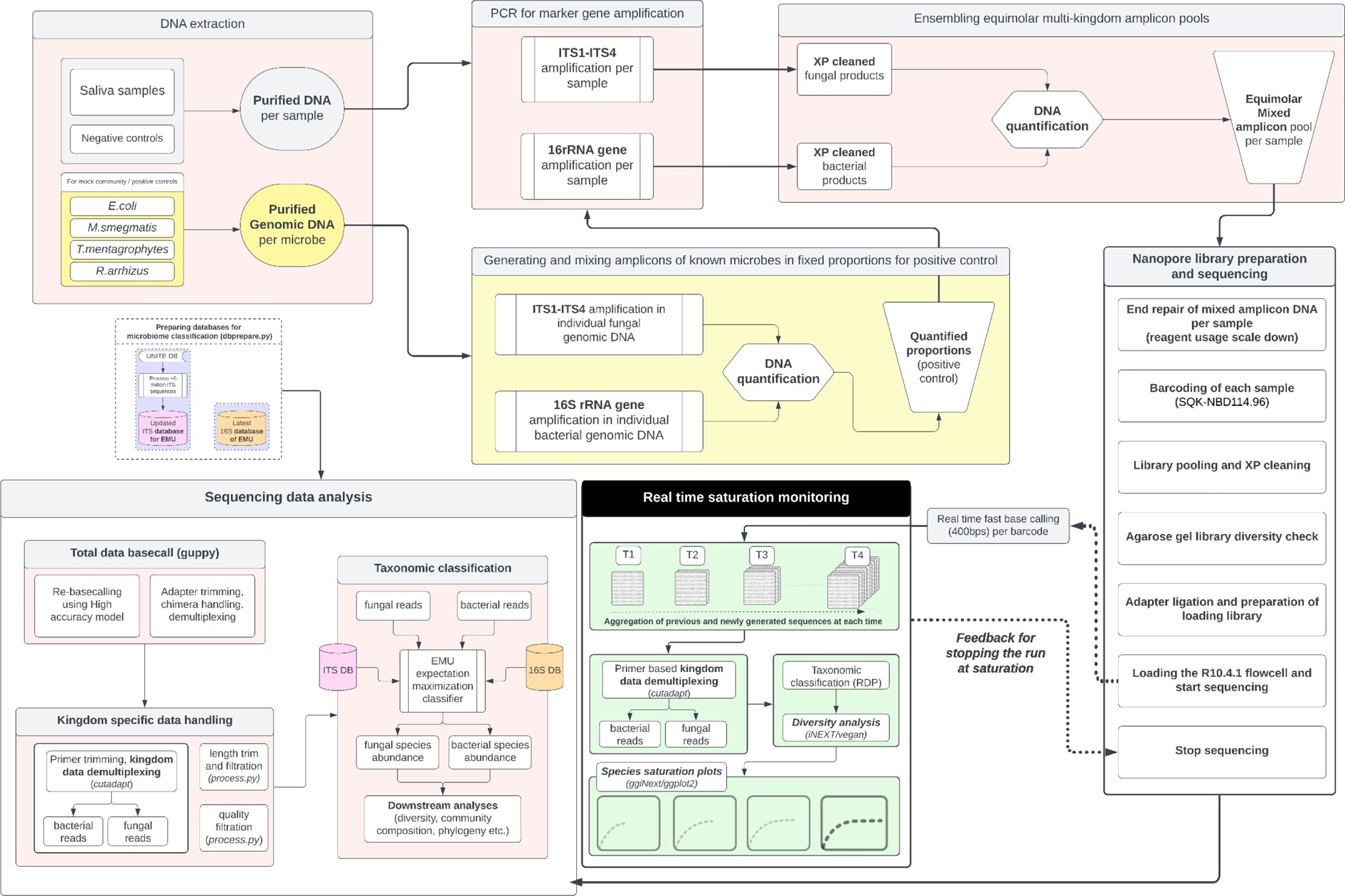
Entire workflow of Ensembleseq providing schematic details of the various steps of both wet and dry lab experiments. These include DNA extraction, maintenance of negative and positive controls, PCR for amplicon generation, ensembling of multi-kingdom amplicons, mixed amplicon library preparation, nanopore sequencing, real time saturation monitoring, database preparation for microbiome classification and sequence data analysis (including species level identification of bacterial and fungal communities.

## 2. Methods

### 2.1. Sample Collection and DNA Extraction

Human saliva samples were collected from 93 subjects with dyspeptic symptoms visiting the Department of Gastroenterology, Post Graduate Institute of Medical Science, Chandigarh, India upon informed consent after due approval of the designated Institutional Human Ethics Committee, in accordance with the guidelines of the Declaration of Helsinki, 1975 was obtained (IEC No. INT/IEC/2021/SP-809). Saliva was collected as a natural drool into 1.5-ml sterile EP tubes, mixed with RNAlater and stored at −80°C.

DNA was extracted using DNeasy Blood & Tissue kit (QIAGEN, Germany) according to the manufacturer’s instructions, with additional steps of bead beating and lysozyme treatment. Briefly, 200 μl saliva sample was diluted in 1000μl tris-EDTA (TE) buffer (pH 8.0) in a 2 ml sterile screw cap vial (Biospec) and centrifuged at 10,000g for 5 min. The pellet was resuspended in 190 μl TE and 10 μl (20 mg/ml) lysozyme and the suspension was incubated at 37°C for 30 min. A mix of 0.1 mm and 1 mm sterile glass beads (Biospec) weighing 100 mg each was added to the vial and homogenization was performed for 1 min using 200 μl disruptor genie lysis buffer and manufacturer’s instructions were followed for subsequent steps. DNA quality was assessed on a 1% w/v agarose gel and quantification was performed using Qubit 3.0. Negative control (reagent only) samples were maintained in each batch of DNA extraction.

### 2.2. PCR amplification of 16S and ITS region

#### 2.2.1. Amplicons for bacteriome

Given the advantage of full-length sequencing using Oxford Nanopore Technology (ONT), entire 16S gene was targeted for amplification using the 27F (AGRGTTYGATYMTGGCTCAG) and 1492R (CGGYTACCTTGTTACGACTT) primers (18). PCR was set up in a set of two technical repeats for each sample as follows: 15 ng template, 0.3 μl (10 μM) 27F primer, 0.3 μl (10 μM) 1492R primer, 0.3 μl 100% DMSO, 7.5 μl LongAmp Taq 2X ReadyMix (NEB) with volume made upto 15 μl with nuclease free water. The polymerase chain reaction consisted of 1 cycle of [denaturation (95 °C,, 5 min)], 35 cycles of [denaturation (95 °C, 60 s), annealing (56 °C, 30 s), extension (65 °C, 75 s)]; 1 cycle of [extension (65 °C, 10 min)].

##### Inclusion of positive and negative control

One set of PCR reactions, as a positive control sample consisting of pre-quantified templates of 16S amplicons specific to routinely used laboratory strains *Mycobacterium smegmatis mc^2^155* and *Escherichia coli DH5α* in 1:4 proportion, was also performed. This was done to facilitate a post-hoc analysis of potential PCR amplification and sequencing bias (if any). PCR was also similarly performed for negative controls. The (technical repeat) reactions were pooled for each sample.

All pools of amplicons were purified using a 1.8X Ampure XP clean-up as per manufacturer’s instructions and eluted in 15μl nuclease free water. Quantification of the cleaned products was performed using Qubit 3.0.

#### 2.2.2. Amplicons for mycobiome

Full length ITS region was targeted through a semi-nested PCR using two sets of primers [39,40] in the following set-up: Outer primer PCR (step 1): 30 ng template, 0.3 μl (10 μM) ITS1F primer (CTTGGTCATTTAGAGGAAGTAA), 0.3 μl (10 μM) ITS4 primer (TCCTCCGCTTATTGATATGC) (40, 41), 0.3 μl 100% DMSO, 5 μl LongAmp Taq 2X readymix with volume made upto 10μl with nuclease free water. The reaction consisted of 1 cycle of [denaturation (95 °C, 5 min)], 35 cycles of [denaturation (95 °C, 60 sec), annealing (56 °C, 30 sec), extension (65 °C, 60 sec)], 1 cycle of extension (65 °C, 10 min).

Inner primer PCR (step 2): 3 μl of step 1 product as template, 0.3 μl (10 μM) ITS1 primer (TCCGTAGGTGAACCTGCGG) (40, 41), 0.3 μl (10 μM) ITS4 primer (TCCTCCGCTTATTGATATGC), 0.3 μl 100% DMSO, 7.5 μl LongAmp Taq 2X readymix with volume made upto 15μl with nuclease free water. The reaction consisted of 1 cycle of [denaturation (95 °C, 5 min)], 15 cycles of [denaturation (95 °C, 60 sec), Annealing (56 °C, 30 sec), extension (65 °C, 60 sec)], 1 cycle of extension (65 °C, 10 min).

##### Positive and negative controls

Similar to bacteriome, a positive control sample consisting of pre-quantified templates of ITS amplicons specific to clinical strains available in the lab, i.e. *Trichophyton interdigitale* UCMS-IGIB-CI1 (GenBank Accession ID: OR899257) and *Rhizopus arrhizus* CAFI-CI-07 (GenBank Accession ID: OR889685), (in 1:2.7 proportion), was also performed. PCR was similarly performed for negative controls. Nested amplicons were purified using a 1.8X Ampure XP cleaning as per manufacturer’s instructions, eluted in 15μl nuclease free water and quantified using Qubit 3.0.

One microlitre from all negative control cleaned products were all pooled together to create a single ‘Negative Control Pool (NCP)’. This can help minimise the need for sequencing multiple negative controls when resources are limited, without neglecting the contaminants across different batches of upstream sample processing. The NCP and one randomly chosen negative control sample were selected for further processing and eventual sequencing. This resulted in a complete plate i.e., a total of 96 samples (93 subjects, 2 negative controls and 1 positive control) for the subsequent workflow.

### 2.3. Native barcoding at scale, ensemble library preparation

ONT currently offers a single specialized-benchmarked kit i.e., 16S barcoding kit (SQK-16S024) for microbiome research, focused only on bacteriome profiling (https://store.nanoporetech.com/16s-barcoding-kit-1-24.html). It allows a maximum of 24 unique samples in a single run using the proprietary barcoded-16S primers, limiting the utility in studies aiming for scale in MINION devices. Given that 16S barcoding has been the default and benchmarked choice, many prior published reports of microbiome research using nanopore sequencing are often observed to be limited by scale in individual runs (15, 18, 42–44). In order to increase the scale of the captured microbiome, we sought to adopt the Native Barcoding Kit – 96 (SQK-NBD114.96) instead to natively barcode the sample specific amplicons (total 96 samples) entailing end-repair/end-prep, PCR free barcoding and adapter ligation. Furthermore, given our goal of concomitantly characterizing both bacteriome and mycobiome of each sample, an ensemble approach was adopted. This method avoids the use of separate library preparation and flow cells for the sequencing of the 16S and ITS amplicons, thereby enabling a cost-effective multi-kingdom interrogation (described in Table 1 and Supplementary file S1):

**Table 1.**
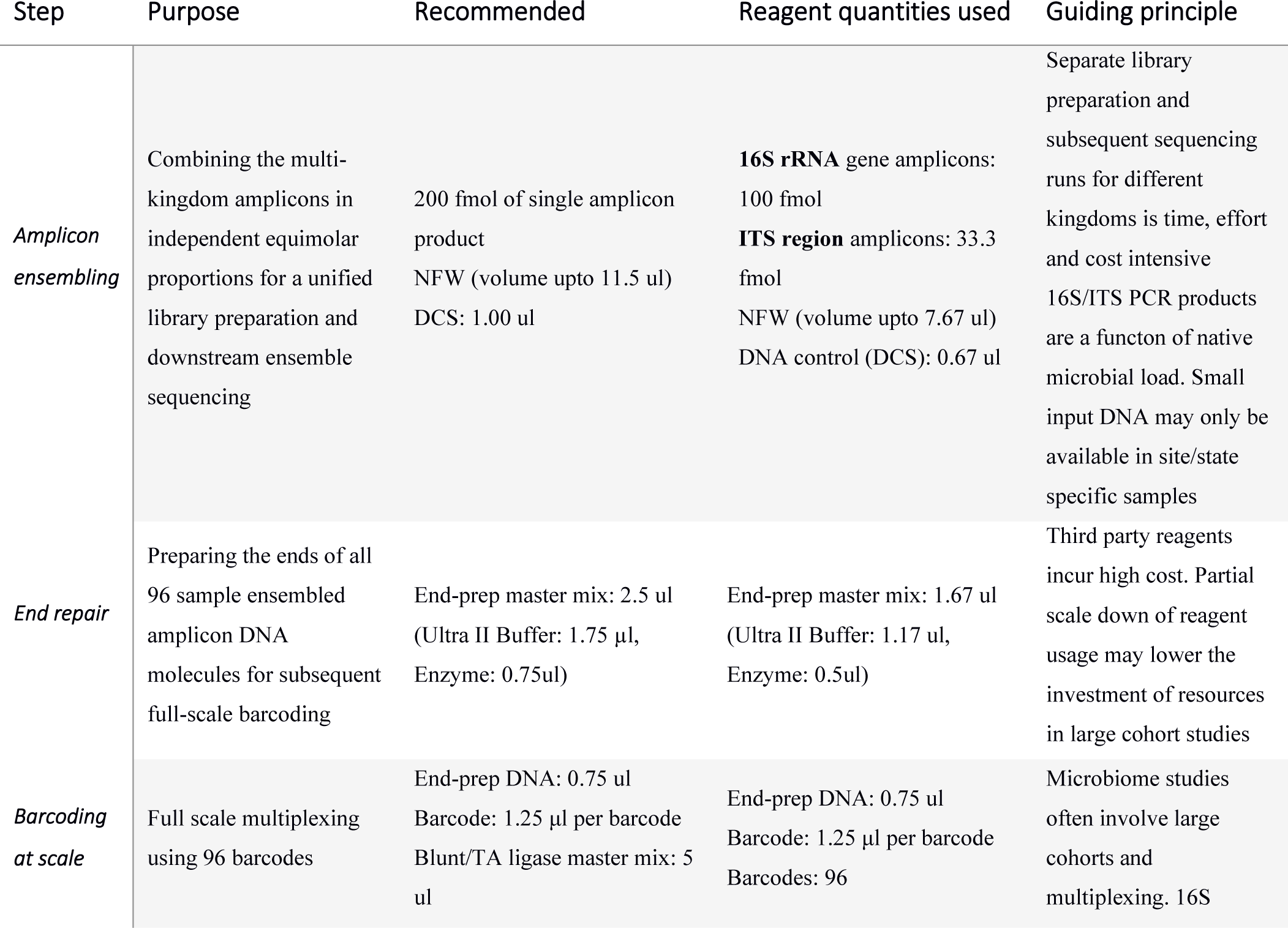
Key steps and guiding principles of EnsembleSeq’s library preparation workflow.

| Step | Purpose | Recommended | Reagent quantities used | Guiding principle |
| --- | --- | --- | --- | --- |
| **Amplicon ensembling** | Combining the multi-kingdom amplicons in independent equimolar proportions for a unified library preparation and downstream ensemble sequencing | 200 fmol of single amplicon product<br>NFW (volume upto 11.5 ul)<br>DCS: 1.00 ul | **16S rRNA** gene amplicons: 100 fmol<br>**ITS region** amplicons: 33.3 fmol<br>NFW (volume upto 7.67 ul)<br>DNA control (DCS): 0.67 ul | Separate library preparation and subsequent sequencing runs for different kingdoms is time, effort and cost intensive<br>16S/ITS PCR products are a function of native microbial load. Small input DNA may only be available in site/state specific samples |
| **End repair** | Preparing the ends of all 96 sample ensembled amplicon DNA molecules for subsequent full-scale barcoding | End-prep master mix: 2.5 ul (Ultra II Buffer: 1.75 µl, Enzyme: 0.75ul) | End-prep master mix: 1.67 ul (Ultra II Buffer: 1.17 ul, Enzyme: 0.5ul) | Third party reagents incur high cost. Partial scale down of reagent usage may lower the investment of resources in large cohort studies |
| **Barcoding at scale** | Full scale multiplexing using 96 barcodes | End-prep DNA: 0.75 ul<br>Barcode: 1.25 µl per barcode<br>Blunt/TA ligase master mix: 5 ul | End-prep DNA: 0.75 ul<br>Barcode: 1.25 µl per barcode<br>Barcodes: 96 | Microbiome studies often involve large cohorts and multiplexing. 16S |

### 2.4. Sequencing and real-time species saturation monitoring

The sequencing library was loaded into an R10.4.1 flow cell (FLO-MIN114) in an Mk1c machine running the MinKNOW version 23.04.5. Fast basecalling (5kHz 400bps) was enabled during sequencing for real time access to the DNA sequences. For each barcode, the passed reads were subjected to real-time taxonomic classification adopting a read accumulation approach (Figure 2 and Figure S1). Real time taxonomic classification was performed for bacteriome and mycobiome using (fast approximate) ribosomal database project (RDP) classifier and species saturation curves were dynamically plotted for each accumulated window using iNEXT and ggplot2 of the R programming language. The sequencing was stopped when sequence generation started plateauing and majority of the samples saturated in improvement for bacterial and fungal taxa identification between 3 pore-scans (especially for the samples with least number of accumulated reads).

### 2.5. High accuracy (re)base calling and sequence data preprocessing

After completion of the sequencing run, the raw signal (pod5) data was rebase-called using a high accuracy (400bps, 5khz, HAC) model of guppy-basecaller on a Windows OS based workstation (RAM: 64gb, NVIDIA RTX A4000, 24 cores of 3.4ghz processor). Sequences were demultiplexed as per the (96) barcode profile of the native barcoding kit and chimeric sequences were disallowed through read splitting during the basecalling. The passed reads were subjected to primer trimming and segregation of bacterial and fungal reads using cutadapt (version 4.4). Reads with minimum read length of 200 and maximum length of 3500 were retained. The quality profile of the passed reads was visualized using Nanoplot (45) and Fastqc (46). Following the quality profile observed in Fastqc for each position across the observed length of reads, read tails were trimmed to a final length of 1550 and a further quality filtration of mean phred 20 was applied on the HAC passed reads using the Fastqc approach.

### 2.6. Taxonomic classification

Taxonomic classification was performed at the CSIR-IGIB high performance computing (HPC) facility using Emu (15). It may also be noted that, while the latest database for bacterial classification was available in Emu (v1.0), the fungal ITS database was outdated. A recent version of fungal database for Emu was therefore created by employing more than 6 million full length ITS sequences obtained from the latest release of UNITE ITS fasta database (47).

#### Data analysis

Species saturation was analysed through rarefaction curve using Vegan library of R programming language (48). Sample level relative abundances of top 10 phyla and species, with highest number of cumulative reads in the study, was observed across all 96 samples (including negative and positive controls). Relative proportion of the top 10 genera was also analyzed. Data visualization was performed using ggplot2 (49) library of R. Cladograms representing the taxonomic lineage of the observed bacterial and fungal population, with at least 0.01% of relative proportion were generated using Graphlan (50).

## 3. Results

### 3.1. EnsembleSeq yields large volume of high-quality data with sufficient species saturation for the combinatorial microbiome in a time-efficient manner

EnsembleSeq approach helped concomitantly capture the major bacteriome as well as the sparse mycobiome in the studied samples without effecting the coverage of the either. Real time monitoring of the species saturation in individual samples was carried out (Figure 3) which indicated the attainment of bacterial as well as the fungal sequence sufficiency for majority of the samples, guiding the timely decision for stopping the run.

**Figure 3.**
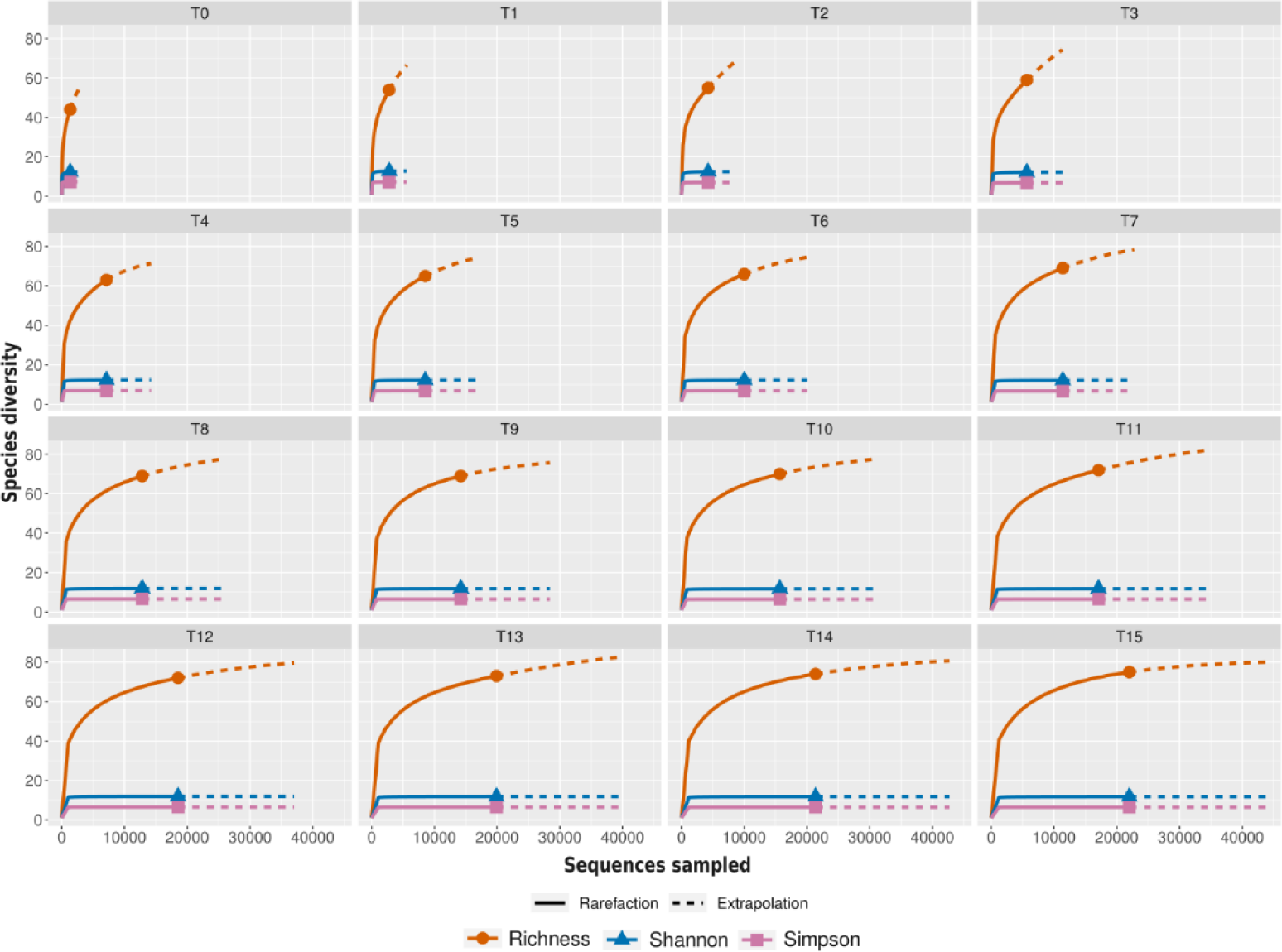
Real time monitoring of bacterial species saturation in the sample specific to barcode1. T0-T15 indicate various time points of real-time sequence data generation with T0 referring to the first period of basecalling after initiating the sequencing. The run was configured to write 4000 reads at a time.

The sequencing run yielded a total population of 7.5 million raw reads with a median read accuracy of 95%. A total of 5.5 million reads passed the high accuracy (HAC) basecalling and 4.9 million (filtered) reads were used for taxonomic classification post length filtration and primer-based segregation of reads. Among these 4.2 million reads pertained to bacteria and 0.7 million pertained to fungi. The proportion of filtered reads (or library size) per sample was approximately uniform (Figure 4a). The length distribution profile of the HAC passed, unfiltered reads exhibited an expected bimodal trend owing to a mix of fungal and bacterial sequences (Figure 4b). Median read length for fungal reads was 556 bases while that of bacterial reads was 1483 bases. The mean read quality for the filtered reads was > Q15 ( > 97% accuracy) for both fungal and bacterial reads as visualized in Nanoplot (Figure 4a) and the modal per basecall quality for each position of the reads was >Q20 ( > 99% accuracy) as visualized using Fastqc (Figure 4a, inset). Final species saturation post high accuracy (HAC) re-basecalling also indicated sufficient capture of both fungal and bacterial diversity (Fig 4c, 4d). Notably the total run time was 30 hours against the recommended default of 72 hours. More than 40% of the nanopores (total 636) remained available for future re-use when the sequencing was stopped.

**Figure 4.**
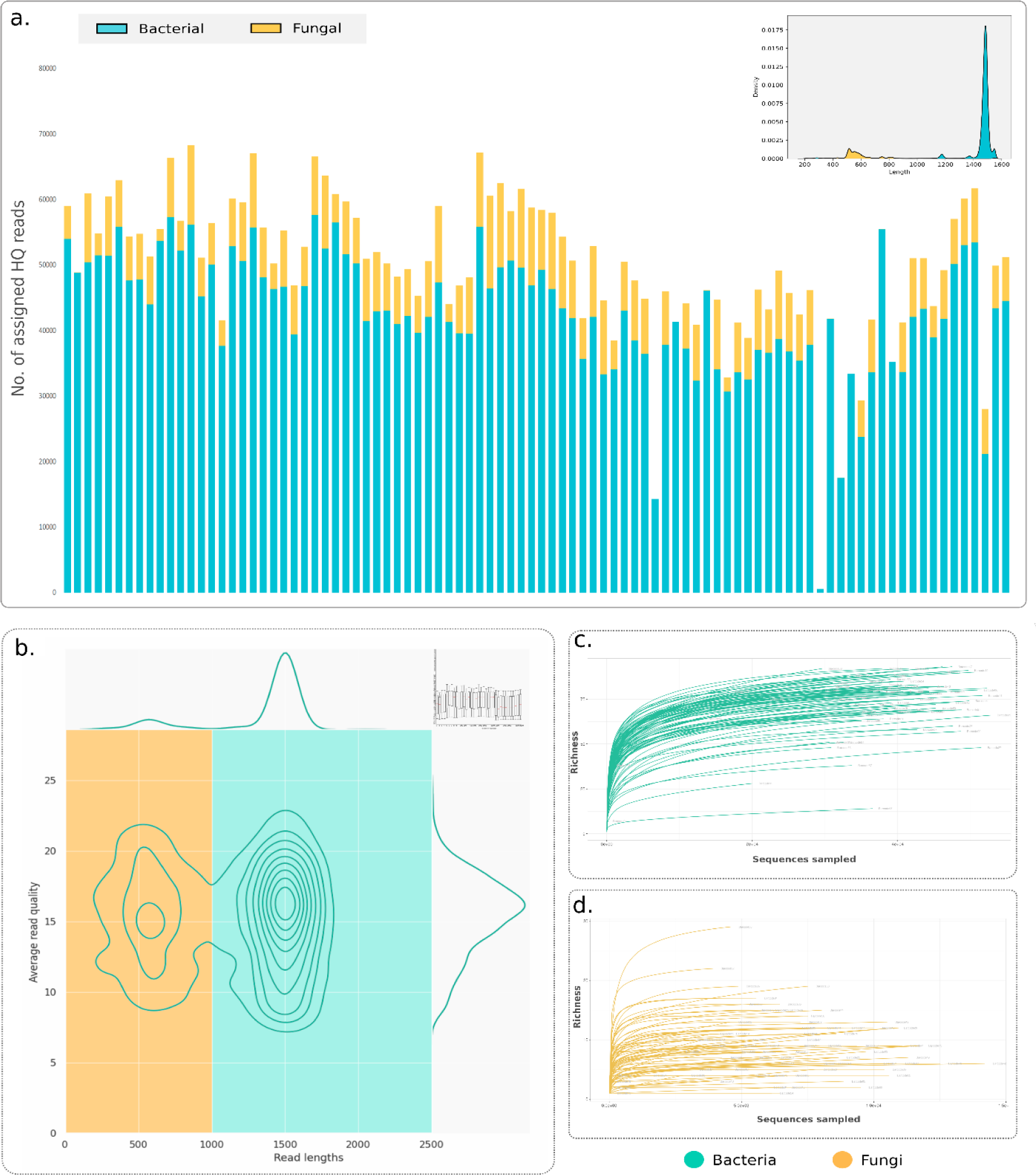
Summary of the data size, quality and species saturation**. (a)** Number of high-quality assigned reads in the samples of this study. One outlier sample (barcode 55 with > 240000 reads) was omitted from this chart. Read length distribution for all the assigned reads in the study is presented in the inset **(b)** Nanoplot generated summary of average read quality (>96% average read accuracy), and in-set Fastqc generated per base position quality trend (>99% modal accuracy per base call) **(c)** Species saturation curves for bacteriome **(d)** Species saturation curves for mycobiome.

### 3.2. Observed species level (bacterial and fungal) taxonomic assignments

#### Taxonomic units of the oral bacteriome

The 4.2 million reads pertaining to bacteria were assigned to 401 species, affiliated to 125 genera and 5 dominant phyla namely Firmicutes, Proteobacteria, Fusobacteria, Bacteroidetes and Actinobacteria (Figure S2a). Five sparse phyla with a total of 5701 reads assigned to them pertained to Spirochaetes (2547 reads), Candidatus Saccharibacteria (2306 reads), Synergistetes (768 reads), Tenericutes (55 reads) and Cynobacteria (25 reads). In total 466 reads remained unassigned at any level of taxonomic hierarchy of bacterial kingdom. The classification results include the assignments for positive and negative controls.

Oral microbiota of the subjects of this study was dominated by the bacteria affiliated to Firmicutes, Proteobacteria and Bacteroidetes - the top three phylum level read assignments observed across all samples (Figure S2a, Figure 5a-d). The top three species level affiliations pertained to *Streptococcus mitis, Neisseria subflava and Streptococcus oralis* observed consistently across the subjects (Figure 5a). At the genus level, 40.7% of the total reads were assigned to *Streptococcus*, followed by *Neisseria* (9.8%), *Veillonella* (9.6%), Prevotella (8.4%), *Granulicatella* (7.2%), *Gemella* (4.6%), *Haemophilus* (2.1%), *Porphyromonas* (2.1%), *Rothia* (1.9%), *Oribacterium* (1.3%) and *Stomatobaculum* (1%), among the top 10 assigned genera (Figure 5c). The cladogram in Figure 5d depicts the phylum and class level affiliation of all the identified bacterial genera in the study with at least 0.01% of assigned reads.

**Figure 5.**
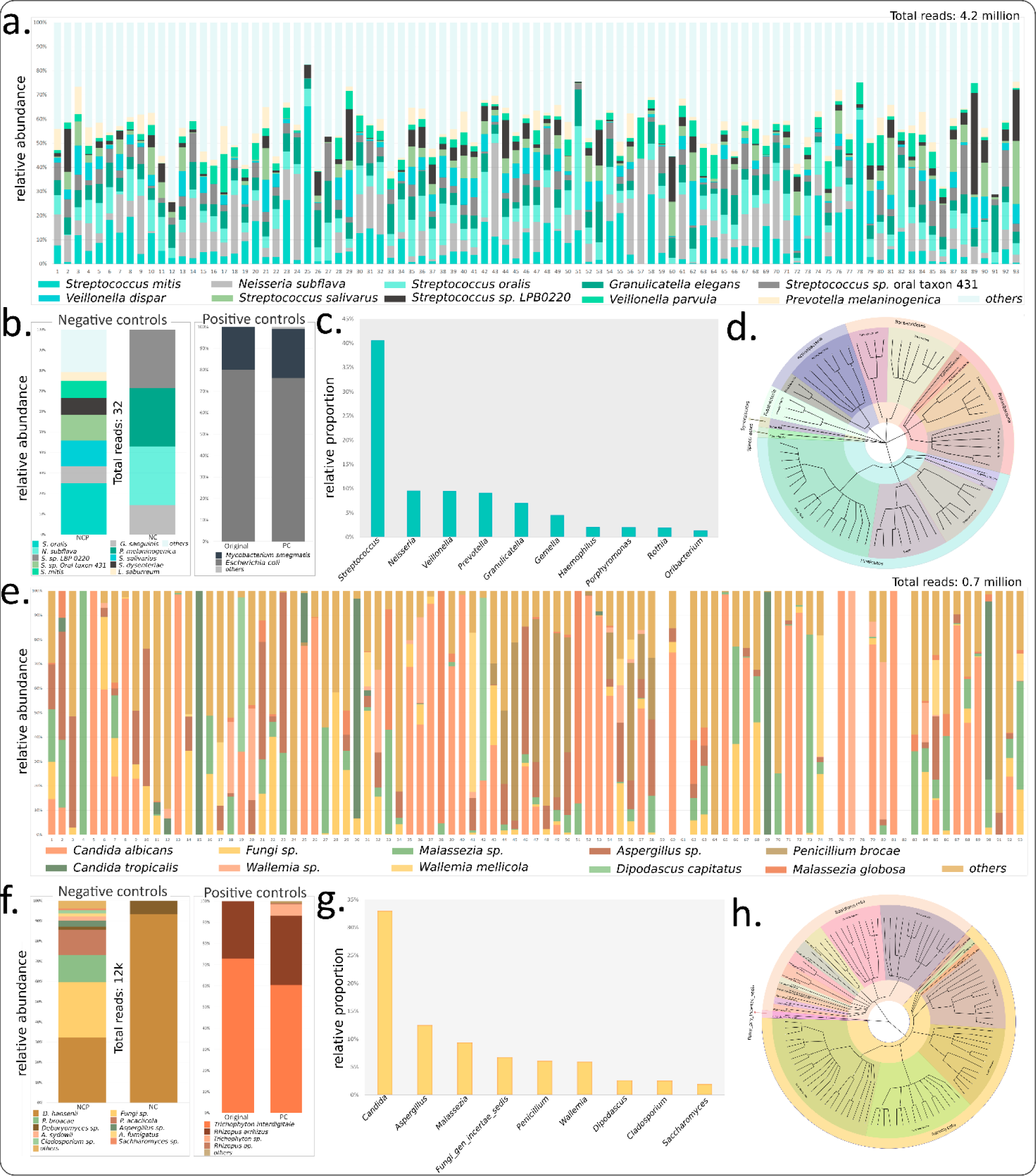
Relative abundance of high-quality assigned reads in the samples, their taxonomic specifications, and the affiliations for control samples of this study. (a) Species level assignments of Top 10 taxonomic units of bacteriome (b) control samples of bacteriome and proportions observed in positive control set (c) top genus level assignment for bacteriome (d) cladogram depicting the taxonomic lineage highlighting phylum and class level taxonomic affiliation of the bacterial genera with at least 0.01% of the assigned reads. Panels 5e-h represent the corresponding results for mycobiome.

#### Top taxonomic units of the oral mycobiome

The ∼0.7 million reads of fungi on the other hand were assigned to 242 species of fungi, pertaining to 130 genera and 4 phyla (Ascomycota, Basidiomycota, Mucormycota and one uncertain phylum) while 5243 reads remained unassigned in total (Figure S2b). This included the assignments for positive and negative controls.

The top two dominant fungal phyla across all samples of the study pertained to Ascomycota and Basidiomycota, along with an uncertain phylum affiliation observed across many samples (Figure S2b, Figure 5e-h). *Candida albicans,* a frequently reported fungal member of the oral microbiota, was observed to be the dominant fungal species across majority of the samples, followed by *Malassezia sp., and Aspergillus sp*. among the other species level designations (Figure 5e). Among the total fungal reads, a majority (33.0%) were assigned to the *Candida* genus, followed by *Aspergillus* (12.6%), *Malassezia* (9.4%), Uncertain genus (7.0%), *Penicillium* (6.2%), *Wallemia* (5.9%), *Dipodascus* (2.6%), *Cladosporium* (2.5%), *Saccharomyces* (1.9%) and *Buckleyzyma* (1.8%) as the top 10 genera with the most assigned reads (Figure 5g). The cladogram in Figure 5h depicts the phylum and class level affiliation of all the identified fungal genera in the study with atleast 0.01% of assigned reads, indicating that Mucormycota is not a key fungal phylum of the oral microbiota. The observed community composition of the bacteria and fungi conformed well with the existing evidence pertaining to the human oral microbiota, as detailed in the Discussion section.

### 3.2. Analysis of the controls indicated minimal contamination with good concordance of species proportions in positive controls

Positive (PC) and negative (NC) control samples were included to assess any bias in amplification, sequencing and assignment. Sufficient saturation was obtained for both the bacteria and fungal PC. The obtained reads were subjected to the defined workflow for confirming species level classification, as carried out for the oral microbiome samples. 34718 and 10423 reads could be assigned respectively to *E.coli* and *M. smegmatis* correctly upto the species level, indicating a close species level conformity with the originally input proportion (1:4) for *E.coli and M. smegmatis* (Figure 5b).

Similar to bacterial, a close conformity between the relative proportions of the originally input proportion of *Trichophyton mentagrophytes and Rhizopus arrhizus* and the observed fungal species proportions in ensemble sequencing of the positive controls (Figure 5f). This indicated the limited bias due to PCR, library preparation, sequencing or taxonomic classification in the overall workflow.

Negative control samples were primarily observed to be devoid of bacterial contamination (Figure 5b). The fungal reads in these controls primarily pertained to *Debaromyces hansenii*, which was not found as a dominant native member of the oral microbiota samples (Figure 5f). Sparse content of *Penicillium brocae, Aspergillus sp*., *Uncertain fungus and Cladosporium sp* reads was also present in the negative controls (Figure 5f).

## 4. Discussion

Multi-kingdom-amplicon sequencing, especially using the state of the art (long read, real-time) Oxford Nanopore Technology, holds promise in low-cost, holistic and species level resolution of the (human) microbiota. Here we attempted a proof-of-concept in concomitantly profiling the oral human bacteriome and mycobiome starting right from the library preparation to a fully multiplexed (96 sample) single sequencing run using MinION (ONT).

In the absence of prior guiding reports for long-read multi-kingdom amplicon sequencing, the key priors (e.g., ideal proportions of mixed amplicons) in our study were guided by the inherent sparsity of the non-bacteriome populations and the need for sufficient coverage for at least the highly abundant and diverse bacterial microflora. We, hence, sought to spike a minor fraction of the full length ITS region (fungal) amplicons (∼33 fmol) to the major proportion of the full length 16S rRNA gene (bacterial) amplicons (∼100 fmol). The said proportions of amplicons were mixed with an anticipation that the minor fraction shouldn’t be too low for the sparsely abundant fungi and the remaining fraction shouldn’t undercover the native bacterial community intended to be captured through the full length 16S rRNA gene sequencing. The choice of priors was supported by the total as well as real time monitoring of the rarefaction which indicated sufficient species saturation for both the fungal and as well as bacterial populations across majority of the samples (Figure. 3 and 4; Figure S1). The latter also enabled timely decision making for stopping the run thereby preserving the flowcell for potential future (re)use. Less than 60% usage of the pores during the entire run (∼ 1 day) of sequencing, covering both bacteriome and mycobiome in the ∼7 million raw and ∼5 million high quality reads at a fully multiplexed (96 sample) scale was encouraging.

Analysis of the obtained bacteriome identified *Streptococcus mitis,* a normal commensal of human oral cavity (51), as the most dominant bacterial species in the entire population of reads in this study. Previously, microbiome reports, albeit at genus level resolution, have consistently reported *Streptococcus* spp. as the primary bacterial genus in the human saliva and oropharynx samples, with *Veilonella* spp*., Neisseria* spp*., Prevotella* spp*., Gemella* spp.*, Granulicatella* spp.*, Porphyromonas spp.* being the other frequently reported genera of the oral cavity (8, 33, 52, 53). Consistent with these previous reports, the current study could identify, at a species level resolution, *Neisseria subflava, Streptococcus oralis, Veilonella parvula, Veilonella dispar, Granulicatella elegans, Streptococcus salivarus, Prevotella melaninogenica* as the other dominant species (Fig. 5) following *S. mitis*. These observations, even though specific to Indian subjects, also exhibit a good concordance with the recently characterized core oral microbiota of European cohort through cost and data intensive whole-genome sequencing (54). The WGS based taxonomic classification by Caselli and colleagues (54) had identified *Streptococcus mitis* as the most abundant bacterial species in the oral cavity along with *Streptococcus oralis, Haemophilus parainfluenzae, Prevotella melaninogenica and Neisseria subflava* as other key microbes of the oral microbiome. This further provides encouraging evidence towards suitability of achieving species level resolution for human bacteriome, in concomitance with the mycobiome, in a cost-effective manner through EnsembleSeq.

Simultaneous sequencing of fungal barcoding DNA (ITS region) enabled potential for useful insights into the mycobiota co-inhabiting the observed bacterial members of the oral microbiome. Among the key fungal taxa, species of *Candida* like *C. albicans and C. tropicalis* were consistently observed in this study*. Penicillium spp., Wallemia spp.* along with genera level assignments for *Aspergillus spp., Cladosporium spp. and Saccharomyces spp.,* were also the major contributors to the total read population of 0.7 million (Figure 5). While Candida species (including *C. albicans*) are regarded as the most common oral cavity-colonising fungi (14, 36, 55), an oral mycobiota study from the United States (14) has shown that apart from Candida, *Malassezia* sp. is another dominant commensal member of the oral mycobiome along with *Aspergilllus, Cladosporium, Cryptococcus, Saccharomyces* and more. Furthermore, as observed for the bacteriome, the top fungal species (*C. albicans*) identified through the current workflow had good concordance with the only species level fungal taxon of *C. albicans* reported through WGS by Caselli *et al.* (54). This highlights the utility of amplicon sequencing in sufficient and rather deep capture of the sparsely abundant members of the rare biosphere (in this case fungi), which could be missed by a data demanding WGS. Members of *Penicillium and Wallemia* have also previously been known to be observed in a handful of human oral microbiome (14, 20, 33, 55). Given the understudied and under characterized nature of the human mycobiota due to the limited available reports, along with a relatively nascent stage of fungal classification-databases and algorithms, it is expected that new members may further be identified through future technical advancements.

Use of negative control in this study (including pooled negative controls) helped in verifying the minimal overlap between the core oral mycobiota and the fungal species identified in the negative controls, consolidating the reliability of the observations. Moreover, negative control samples had minimal (total 32 reads) contamination of bacterial reads (Figure 5b, f). It was also encouraging to observe the conformity between the proportions of fungal and bacterial species in positive controls and those obtained in terms of the sequenced reads, indicating the potential of accurate species identification as well as controlling and monitoring the factors that can contribute to the bias in community profiling. It is hoped that our case study and observations, hereby presented as a step-by-step outlined EnsembleSeq workflow, may guide the future attempts at further optimizing the right proportion of input molecules, scope of further scale-up and attempting multiple amplicons (beyond bacteriome and mycobiome) without compromising the expected saturations. We further summarize potential improvements and future directions as the following:

i. **Lower proportions of fungal spike and going beyond bacteriome and mycobiome** Success in sufficiently capturing the sample specific mycobiota and bacterial species through a small (25%) spike of fungal ITS DNA in the total library was observed in the current study. This opens the avenue for attempting/optimizing further lower proportions, which may guide the scope for simultaneous spiking of multiple kingdom specific barcoding DNA at the same time.
ii. **Multiplexing 16S and ITS region amplification in a single PCR run** In the current research, separate amplification of 16S and ITS region amplification was carried out for all 96 samples. PCR multiplexing is however also theoretically possible and attempts at optimizing the template amount, primer concentrations, annealing conditions may yield success thereby further reducing the time and cost associated with multi-amplicon sequencing based microbiome studies. While this may limit the control over equimolar libraries, post-hoc normalizations can potentially address the skewed library sizes.
iii. **Towards adaptive barcode prioritization for amplicon (nanopore) sequencing** Real-time access to the DNA sequences enabled continuous monitoring of the species saturation through dynamic classification and rarefaction analysis of the generated reads. With the availability of adaptive sequencing API, this utility of dynamic saturation monitoring may further be extended to barcode prioritization, such that the barcodes that have already saturated for all target microorganisms can be omitted for further sequencing, thereby further optimizing the usage of the sequencing resources.

## 5. Conclusion

With the advent of high accuracy long-read sequencing methods, an interest has been ignited towards relooking at the human microbiota at a higher resolution. At this juncture, it would be prudent to take strides beyond the traditional approach of treating bacteriome as the “microbiome” and conduct experiments that aim to simultaneously profile different microbial kingdoms of the studied samples. Amplicon sequencing, given its cost effectiveness and ease of data handling, is preferred by microbiome scientists across the world for initial insights and taxonomic characterization of microbial communities. EnsembleSeq workflow presented in this proof-of-concept report, was attempted with the goal of targeting multiple kingdoms of microorganisms, specifically bacteriome and mycobiome of the total human oral microbiome in a resource and time efficient manner. The observations indicated the potential to simultaneously identify both fungal and bacteria species, without compromising on coverage, resolution, or the native community composition, in a single fully multiplexed (96 sample) nanopore amplicon sequencing run. Dynamic tracking of species saturation, enabled by real time availability of the sequenced DNA, further opens avenues for optimizing the time of run (and even real time barcode prioritization). Guiding studies are currently needed to add to the confidence of microbiome scientific community that holistic and species level snapshots of human microbiome can be achieved without incurring significant costs of time, effort or resources. We hope to have re-ignited further thought in this direction.

## Acknowledgements

Authors would like to thank the Director CSIR-IGIB and CSIR for the financial support towards executing this research. Authors would also like to acknowledge the HPC facility at CSIR-IGIB in continuous management of the compute resources. SN would like to thank the doctoral advisory committee (DAC) and lab members for their valuable suggestions and invigorating discussions.

## Authors’ Contribution

**Conceived idea, Method design**: SN, BT

**Sample transport, DNA purification, PCR, library prep, sequencing, analysis, figures:** SN

**Subject recruitment, Sample collection:** HH, UD

**Manuscript writing and editing**: SN, BT, SSM

**Supervision**: BT, SSM.

## References

1. Ursell LK, Metcalf JL, Parfrey LW, Knight R. 2012. Defining the human microbiome. Nutr Rev 70.

2. Lane N. 2015. The unseen World: Reflections on Leeuwenhoek (1677) ‘Concerning little animals.’ Philosophical Transactions of the Royal Society B: Biological Sciences 10.1098/rstb.2014.0344.

3. Baker JL, Bor B, Agnello M, Shi W, He X. 2017. Ecology of the Oral Microbiome: Beyond Bacteria. Trends Microbiol 10.1016/j.tim.2016.12.012.

4. Vemuri R, Shankar EM, Chieppa M, Eri R, Kavanagh K. 2020. Beyond just bacteria: Functional biomes in the gut ecosystem including virome, mycobiome, archaeome and helminths. Microorganisms 10.3390/microorganisms8040483.

5. Monteiro-da-Silva F, Sampaio-Maia B, Pereira M de L, Araujo R. 2013. Characterization of the oral fungal microbiota in smokers and non-smokers. Eur J Oral Sci 121.

6. Narunsky-Haziza L, Sepich-Poore GD, Livyatan I, Asraf O, Martino C, Nejman D, Gavert N, Stajich JE, Amit G, González A, Wandro S, Perry G, Ariel R, Meltser A, Shaffer JP, Zhu Q, Balint-Lahat N, Barshack I, Dadiani M, Gal-Yam EN, Patel SP, Bashan A, Swafford AD, Pilpel Y, Knight R, Straussman R. 2022. Pan-cancer analyses reveal cancer-type-specific fungal ecologies and bacteriome interactions. Cell 185.

7. Huffnagle GB, Noverr MC. 2013. The emerging world of the fungal microbiome. Trends Microbiol 10.1016/j.tim.2013.04.002.

8. Tu Y, Zhou Z, Shu C, Zhou Y, Zhou X. 2022. The Crosstalk Between Saliva Bacteria and Fungi in Early Childhood Caries. Front Cell Infect Microbiol 12.

9. Grant KR. 2022. Next-Generation Amplicon Sequencing: A Cost-Effective Method for Exploring Microbial BiodiversityMolecular Genetics and Genomics Tools in Biodiversity Conservation.

10. Meslier V, Quinquis B, Da Silva K, Plaza Oñate F, Pons N, Roume H, Podar M, Almeida M. 2022. Benchmarking second and third-generation sequencing platforms for microbial metagenomics. Sci Data 9.

11. Clarridge JE. 2004. Impact of 16S rRNA gene sequence analysis for identification of bacteria on clinical microbiology and infectious diseases. Clin Microbiol Rev 10.1128/CMR.17.4.840-862.2004.

12. Schriefer AE, Cliften PF, Hibberd MC, Sawyer C, Brown-Kennerly V, Burcea L, Klotz E, Crosby SD, Gordon JI, Head RD. 2018. A multi-amplicon 16S rRNA sequencing and analysis method for improved taxonomic profiling of bacterial communities. J Microbiol Methods 154.

13. Nilsson RH, Ryberg M, Abarenkov K, Sjökvist E, Kristiansson E. 2009. The ITS region as a target for characterization of fungal communities using emerging sequencing technologies. FEMS Microbiol Lett 296.

14. Dupuy AK, David MS, Li L, Heider TN, Peterson JD, Montano EA, Dongari-Bagtzoglou A, Diaz PI, Strausbaugh LD. 2014. Redefining the human oral mycobiome with improved practices in amplicon-based taxonomy: Discovery of Malassezia as a prominent commensal. PLoS One 9.

15. Curry KD, Wang Q, Nute MG, Tyshaieva A, Reeves E, Soriano S, Graeber E, Finzer P, Mendling W, Wu Q, Savidge T, Villapol S, Dilthey A, Treangen TJ. 2021. Emu: Species-Level Microbial Community Profiling for Full-Length Nanopore 16S Reads. bioRxiv.

16. Szoboszlay M, Schramm L, Pinzauti D, Scerri J, Sandionigi A, Biazzo M. 2023. Nanopore Is Preferable over Illumina for 16S Amplicon Sequencing of the Gut Microbiota When Species-Level Taxonomic Classification, Accurate Estimation of Richness, or Focus on Rare Taxa Is Required. Microorganisms 11.

17. Johnson JS, Spakowicz DJ, Hong BY, Petersen LM, Demkowicz P, Chen L, Leopold SR, Hanson BM, Agresta HO, Gerstein M, Sodergren E, Weinstock GM. 2019. Evaluation of 16S rRNA gene sequencing for species and strain-level microbiome analysis. Nat Commun 10.

18. Matsuo Y, Komiya S, Yasumizu Y, Yasuoka Y, Mizushima K, Takagi T, Kryukov K, Fukuda A, Morimoto Y, Naito Y, Okada H, Bono H, Nakagawa S, Hirota K. 2021. Full-length 16S rRNA gene amplicon analysis of human gut microbiota using MinION^TM^ nanopore sequencing confers species-level resolution. BMC Microbiol 21.

19. Clavel T, Horz HP, Segata N, Vehreschild M. 2022. Next steps after 15 stimulating years of human gut microbiome research. Microb Biotechnol 10.1111/1751-7915.13970.

20. Lu J, Zhang X, Zhang X, Wang L, Zhao R, Liu XY, Liu X, Zhuang W, Chen L, Cai L, Wang J. 2022. Nanopore sequencing of full rRNA operon improves resolution in mycobiome analysis and reveals high diversity in both human gut and environments. Mol Ecol 10.1111/mec.16534.

21. Pedroso-Roussado C, Guppy F, Bowler L, Inacio J. 2023. Nanopore sequencing of DNA barcodes to unveil the diversity of fungal mock communities. Open Research Europe 3.

22. Ohta A, Nishi K, Hirota K, Matsuo Y. 2023. Using nanopore sequencing to identify fungi from clinical samples with high phylogenetic resolution. Sci Rep 13.

23. Jang Y, Kim S, Kim N, Son H, Ha EJ, Koh EJ, Phi JH, Park CK, Kim JE, Kim SK, Lee SK, Cho WS, Moon J, Chu K. 2022. Nanopore 16S sequencing enhances the detection of bacterial meningitis after neurosurgery. Ann Clin Transl Neurol 9.

24. Low L, Fuentes-Utrilla P, Hodson J, O’Neil JD, Rossiter AE, Begum G, Suleiman K, Murray PI, Wallace GR, Loman NJ, Rauz S. 2021. Evaluation of full-length nanopore 16S sequencing for detection of pathogens in microbial keratitis. PeerJ 9.

25. Chen Y, Mao L, Lai D, Xu W, Zhang Y, Wu S, Yang D, Zhao S, Liu Z, Xiao Y, Tang Y, Meng X, Wang M, Shi J, Chen Q, Shu Q. 2023. Improved targeting of the 16S rDNA nanopore sequencing method enables rapid pathogen identification in bacterial pneumonia in children. Front Cell Infect Microbiol 12.

26. Rozas M, Brillet F, Callewaert C, Paetzold B. 2022. MinION^TM^ Nanopore Sequencing of Skin Microbiome 16S and 16S-23S rRNA Gene Amplicons. Front Cell Infect Microbiol 11.

27. Oberle A, Urban L, Falch-Leis S, Ennemoser C, Nagai Y, Ashikawa K, Ulm PA, Hengstschläger M, Feichtinger M. 2021. 16S rRNA long-read nanopore sequencing is feasible and reliable for endometrial microbiome analysis. Reprod Biomed Online 42.

28. Omi M, Matsuo Y, Araki-Sasaki K, Oba S, Yamada H, Hirota K, Takahashi K. 2022. 16S rRNA nanopore sequencing for the diagnosis of ocular infection: a feasibility study. BMJ Open Ophthalmol 7.

29. Rausch P, Rühlemann M, Hermes BM, Doms S, Dagan T, Dierking K, Domin H, Fraune S, Von Frieling J, Hentschel U, Heinsen FA, Höppner M, Jahn MT, Jaspers C, Kissoyan KAB, Langfeldt D, Rehman A, Reusch TBH, Roeder T, Schmitz RA, Schulenburg H, Soluch R, Sommer F, Stukenbrock E, Weiland-Bräuer N, Rosenstiel P, Franke A, Bosch T, Baines JF. 2019. Comparative analysis of amplicon and metagenomic sequencing methods reveals key features in the evolution of animal metaorganisms. Microbiome 7.

30. Brumfield KD, Huq A, Colwell RR, Olds JL, Leddy MB. 2020. Microbial resolution of whole genome shotgun and 16S amplicon metagenomic sequencing using publicly available NEON data. PLoS One 15.

31. Angebault C, Payen M, Woerther PL, Rodriguez C, Botterel F. 2020. Combined bacterial and fungal targeted amplicon sequencing of respiratory samples: Does the DNA extraction method matter? PLoS One 15.

32. Simon GL, Gorbach SL. 1984. Intestinal flora in health and disease. Gastroenterology 86.

33. Heng W, Wang W, Dai T, Jiang P, Lu Y, Li R, Zhang M, Xie R, Zhou Y, Zhao M, Duan N, Ye Z, Yan F, Wang X. 2022. Oral Bacteriome and Mycobiome across Stages of Oral Carcinogenesis. Microbiol Spectr 10.

34. Song Y, Kim MS, Chung J, Na HS. 2023. Simultaneous Analysis of Bacterial and Fungal Communities in Oral Samples from Intubated Patients in Intensive Care Unit. Diagnostics 13.

35. Morrison GA, Fu J, Lee GC, Wiederhold NP, Cañete-Gibas CF, Bunnik EM, Wickes BL. 2020. Nanopore sequencing of the fungal intergenic spacer sequence as a potential rapid diagnostic assay. J Clin Microbiol 58.

36. Patel M. 2022. Oral Cavity and Candida albicans: Colonisation to the Development of Infection. Pathogens 10.3390/pathogens11030335.

37. Kohi S, Macgregor-Das A, Dbouk M, Yoshida T, Chuidian M, Abe T, Borges M, Lennon AM, Shin EJ, Canto MI, Goggins M. 2021. Alterations in the Duodenal Fluid Microbiome of Patients With Pancreatic Cancer. Clinical Gastroenterology and Hepatology 10.1016/j.cgh.2020.11.006.

38. Shay E, Sangwan N, Padmanabhan R, Lundy S, Burkey B, Eng C. 2020. Bacteriome and mycobiome and bacteriome-mycobiome interactions in head and neck squamous cell carcinoma. Oncotarget 11.

39. Hu J, Tang J, Zhang X, Yang K, Zhong A, Yang Q, Liu Y, Li Y, Zhang T. 2023. Landscape in the gallbladder mycobiome and bacteriome of patients undergoing cholelithiasis with chronic cholecystitis. Front Microbiol 14.

40. Langsiri N, Worasilchai N, Irinyi L, Jenjaroenpun P, Wongsurawat T, Luangsa-ard JJ, Meyer W, Chindamporn A. 2023. Targeted sequencing analysis pipeline for species identification of human pathogenic fungi using long-read nanopore sequencing. IMA Fungus 14.

41. Op De Beeck M, Lievens B, Busschaert P, Declerck S, Vangronsveld J, Colpaert J V. 2014. Comparison and validation of some ITS primer pairs useful for fungal metabarcoding studies. PLoS One 9.

42. Waechter C, Fehse L, Welzel M, Heider D, Babalija L, Cheko J, Mueller J, Pöling J, Braun T, Pankuweit S, Weihe E, Kinscherf R, Schieffer B, Luesebrink U, Soufi M, Ruppert V. 2023. Comparative analysis of full-length 16s ribosomal RNA genome sequencing in human fecal samples using primer sets with different degrees of degeneracy. Front Genet 14.

43. Lao HY, Ng TTL, Wong RYL, Wong CST, Lee LK, Wong DSH, Chan CTM, Jim SHC, Leung JSL, Lo HWH, Wong ITF, Yau MCY, Lam JYW, Wu AKL, Siu GKH. 2022. The Clinical Utility of Two High-Throughput 16S rRNA Gene Sequencing Workflows for Taxonomic Assignment of Unidentifiable Bacterial Pathogens in Matrix-Assisted Laser Desorption Ionization-Time of Flight Mass Spectrometry. J Clin Microbiol 60.

44. Lee AWT, Chan CTM, Wong LLY, Yip CY, Lui WT, Cheng KC, Leung JSL, Lee LK, Wong ITF, Ng TTL, Lao HY, Siu GKH. 2023. Identification of microbial community in the urban environment: The concordance between conventional culture and nanopore 16S rRNA sequencing. Front Microbiol 14.

45. De Coster W, D’Hert S, Schultz DT, Cruts M, Van Broeckhoven C. 2018. NanoPack: Visualizing and processing long-read sequencing data. Bioinformatics 34.

46. Andrews S. 2010. FastQC: a quality control tool for high throughput sequence data.

47. Abarenkov KZAPTPRIFNRHKU. 2023. Full UNITE+INSD dataset for Fungi. Version 18.07.2023. UNITE Community.

48. Dixon P. 2003. VEGAN, a package of R functions for community ecology. Journal of Vegetation Science 10.1111/j.1654-1103.2003.tb02228.x.

49. Wickham H. 2011. ggplot2. Wiley Interdiscip Rev Comput Stat 3.

50. Asnicar F, Weingart G, Tickle TL, Huttenhower C, Segata N. 2015. Compact graphical representation of phylogenetic data and metadata with GraPhlAn. PeerJ 2015.

51. Engen SA, Rørvik GH, Schreurs O, Blix IJS, Schenck K. 2017. The oral commensal Streptococcus mitis activates the aryl hydrocarbon receptor in human oral epithelial cells. Int J Oral Sci 9.

52. Wang S, Song F, Gu H, Wei X, Zhang K, Zhou Y, Luo H. 2022. Comparative Evaluation of the Salivary and Buccal Mucosal Microbiota by 16S rRNA Sequencing for Forensic Investigations. Front Microbiol 13.

53. Kilian M, Chapple ILC, Hannig M, Marsh PD, Meuric V, Pedersen AML, Tonetti MS, Wade WG, Zaura E. 2016. The oral microbiome - An update for oral healthcare professionals. Br Dent J 221.

54. Caselli E, Fabbri C, D’Accolti M, Soffritti I, Bassi C, Mazzacane S, Franchi M. 2020. Defining the oral microbiome by whole-genome sequencing and resistome analysis: The complexity of the healthy picture. BMC Microbiol 20.

55. Diaz PI, Hong BY, Dupuy AK, Strausbaugh LD. 2017. Mining the oral mycobiome: Methods, components, and meaning. Virulence 10.1080/21505594.2016.1252015.

