## Supplementary file S1 for "EnsembleSeq: A workflow towards real-time, rapid and simultaneous multi-kingdom amplicon sequencing for holistic and cost-effective microbiome research at scale"

**EnsembleSeq: A workflow towards real-time, rapid and simultaneous multi-kingdom amplicon sequencing for holistic and cost effective microbiome research at scale**

Sunil Nagpal[1,2,3], Sharmila S. Mande[3], Harish Hooda[4], Usha Dutta[4], Bhupesh Taneja[1,2*]

1. CSIR-Institute of Genomics and Integrative Biology, New Delhi, India

2. Academy of Scientific and Innovative Research (AcSIR), Ghaziabad, India

3. TCS Research, Tata Consultancy Services Ltd, Pune, India

4. Department of Gastroenterology, Post Graduate Institute of Medical Education & Research, Chandigarh, India

#### EnsembleSeq library preparation steps

1) For each sample, corresponding bacterial and fungal cleaned amplicons were pooled into an approximate 3:1 proportion to make a 133.3 fmol ensembled sample such that it contained 100 fmol of full length 16S rRNA gene (97.4 ng per sample assuming ~1.5kb molecule length) and 33.3 fmol ITS dsDNA molecules (11.9 ng per sample assuming ~550b molecule length). The volume was made upto 7.67 µl using NFW and 0.67 ul of DNA Control Sample (DCS) was added to the pool as available in the SQK-NBD114.96.

2) To each of the ensemble sample, 1.67 µl of End-prep master mix (containing 17.55 µl NEB Ultra II End-prep reaction buffer and 7.5 µl Ultra II End-prep Enzyme mix) was added.

3) All components were thoroughly mixed as per manufacturer's instructions and incubated in a thermal cycler at 20 °C for 7.5 minutes and 65 °C for 7.5 minutes.

4) From each of the end-prepped ensemble DNA, 0.75 µl of sample was taken and mixed with 3 µl of NFW followed by 1.25 µl of unique barcode from the 96 barcode plate as available in the native barcoding kit.

5) The barcodes were ligated to the ensemble DNA using 5 µl of Blunt/TA ligase master mix and incubated at room temperature for 20 minutes.

6) Each ligation reaction was stopped using 1 µl of EDTA and all barcoded samples were subsequently pooled into two LoBind (Eppendorf) tubes of 1.5 ml each. This yielded ~480 µl of pooled reaction in each LoBind tube.

7) AMPure XP beads were then added to the pooled reactions at 1.5x volume (i.e. 720 µl) and a standard clean-up was performed as per manufacturer's instructions (washing twice with 80% molecular grade ethanol). Elution was peformed in 17.5 µl NFW from each tube, totalling 35 µl of total barcoded ensemble-DNA library.

8) Adapters were ligated to the ensemble-DNA library as instructed in the native barcoding kit:

| **Reagent** | **Volume** |
|---|---|
| Pooled barcoded sample | 30 µl |
| Native Adapter (NA) | 5 µl |
| NEBNext Quick Ligation Reaction Buffer (5X) | 10 µl |
| Quick T4 DNA Ligase | 5 µl |
| **Total** | **50 µl** |

9) Final round of AMPure XP bead based cleaning of adapter ligated library was performed. Instead of alcohol (80% Molecular grade ethanol) washing of beads, this step, as recommended in the native barcoding kit must use 125 µl Short Fragment Buffer (SFB) to avoid downstream damage to the nanopores due to ethanol residues. Additionally unlike magnetic rack mounted gentle wash, beads should be flicked to resuspend, followed by a spin down, and reracking to the magnetic rack to allow the beads to pellet. Supernatant is

subsequently removed using a pipette and stored in a back-up vial for contingency (notably, supernatant can be discarded as recommended by the kit, authors prefered exercising caution and not discarding any supernatant or beads in any of the wash steps of entire protocol to account for potential human errors and hence allowing a chance for recovery of library in case of any contingency). Washing is performed twice as instructed. Elution was performed by resuspending the washed pellet in 15 µl Elution Buffer (EB).

10) Washed, adapter ligated ensemble-DNA library was quantified using Qubit 3.0. 20ng of this library was used to make the final sequencing library:

| Reagent | Volume per flow cell |
|---|---|
| Sequencing Buffer (SB) | 37.5 µl |
| Library Beads (LIB) mixed immediately before use | 25.5 µl |
| DNA library | 12 µl (containing 20ng in EB) |
| **Total** | **75 µl** |

11) The above final library was loaded into the primed R10.4.1 flow cell (FLO-MIN114) mounted in an mk1c machine containing the MinKNOW version 23.04.5.

Reader is advised to refer for further details, the default instructions of the Ligation sequencing amplicons - Native Barcoding Kit 96 V14 (SQK-NBD114.96) protocol.

### Figure S1

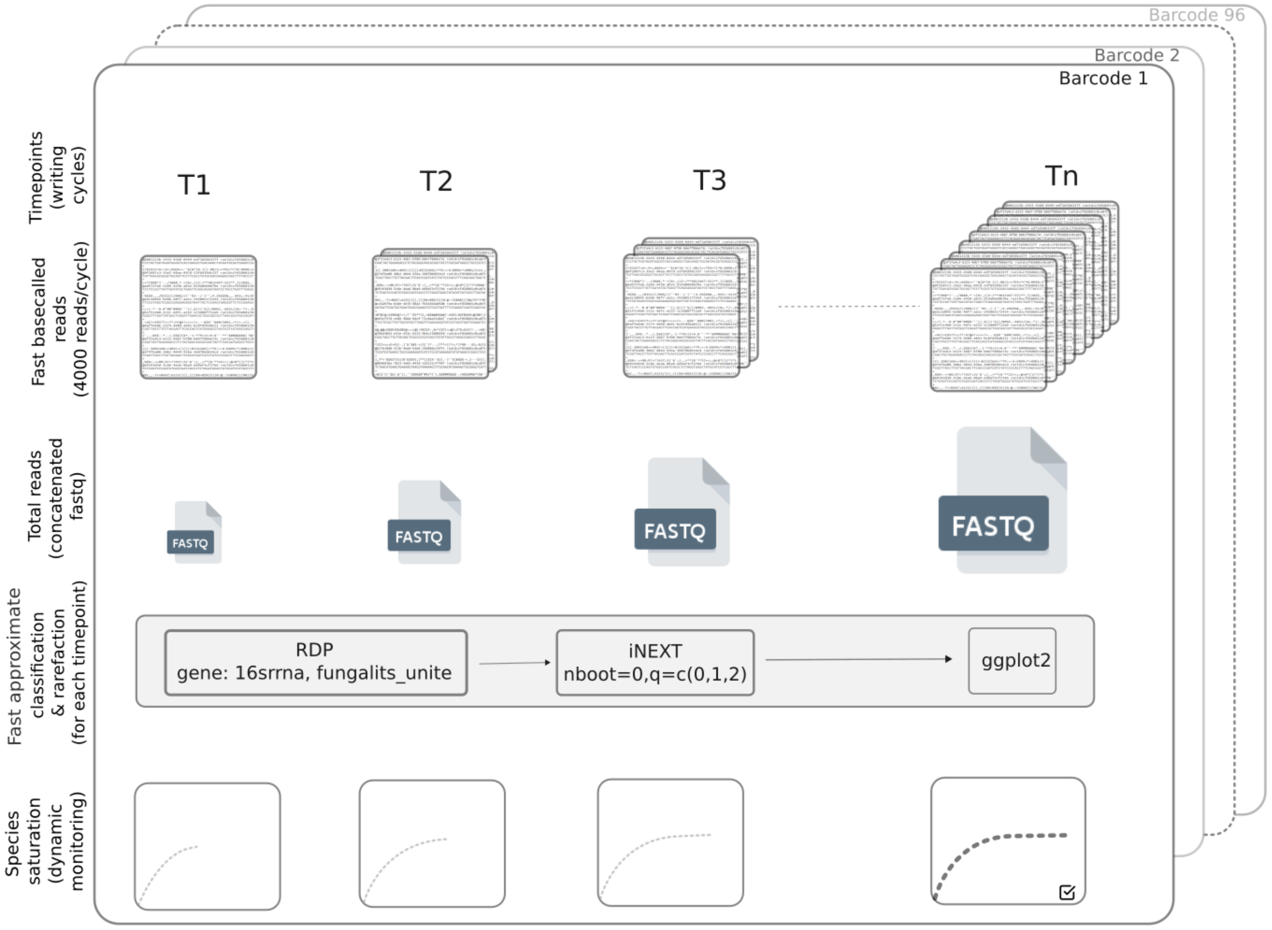


**Figure S1.** Schematic representation of the dynamic monitoring of species saturation that can guide the optimal run-time for nanopore sequencing of ensembled amplicons. T1,T2,T3..Tn represent various time points after starting the sequencing run, where T1 refers to the time when first set of sequences are real-time base called and written on the disc. Subsequent time points refer to the events of new sequences being written on disc (for e.g. nanopore guppy basecaller is configured to write 4000 reads in each writing cycle by default). For each time point, all available reads can be concatenated to a single fastq file and the reads can be subjected to direct taxonomic classification (fast but approximate) using RDP classifier, employing both bacterial and fungal databases. The accumulated classified sequences can then be separately analysed for species saturation through rarefaction analysis (using iNEXT library of R programming language) and the rarefaction plot for the given time point can be plotted using ggplot2 library of R.  Taxonomic classifications of independent time point specific sequences (without sequence accumulation) can also be aggregated to the accumulated taxonomic composition of the previous time point to yied the similar temporal trend of rarefaction. The dynamic monitoring step of EnsembleSeq workflow is coined Raredynamics.

### Figure S2

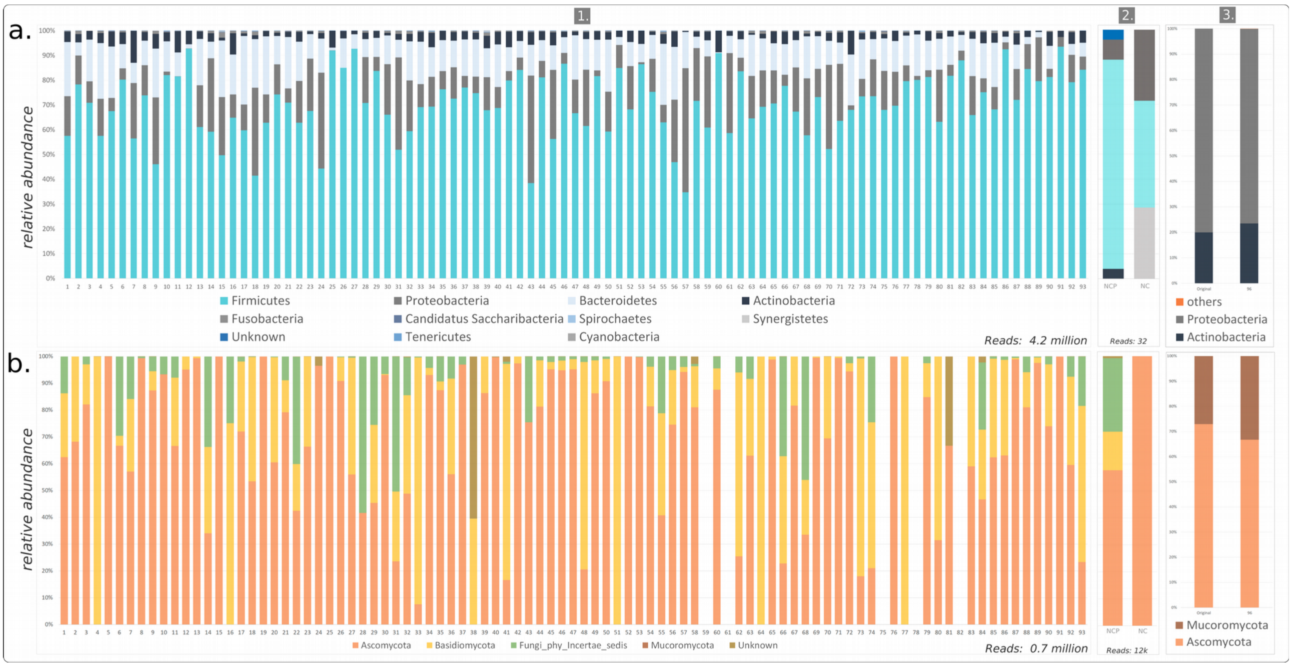


**Figure S2**. Relative abundance of high-quality assigned reads in the samples of this study at phylum level of taxonomic hierarchy for (a) bacteriome and (b) mycobiome. Assignment for negative control samples (NCP and NC) are provided in a2 and b2 while for positive control samples is provided in a3 and b3. Original intended proportion of the positive control molecules are provided in the left stack bar of a3 and b3 panels.
